# Predicting Antibody Self-Association with Sequence–Structure Fusion Models: The Central Role of CSI-BLI in Early Developability Screening

**DOI:** 10.64898/2026.04.13.718222

**Authors:** Shafayat Ahmed, Federico Devalle, Lauren Leisen, Tony Pham, Bismark Amofah, Amber Lee, Mark Hutchinson, Chacko Chakiath, Jen DiChiara, Sharfa Farzandh, Madi Kreitz, Alison Hinton, Neil Mody, Andrew Dippel, Gilad Kaplan, Maryam Pouryahya

## Abstract

Antibody-based biologics are expanding rapidly, yet challenges in development from self-association, high viscosity, aggregation, and unfavorable clearance underscore the need for accurate in silico screening. Clone self-interaction biolayer interferometry (CSI-BLI) is a plate-based, low-material assay of weak, reversible self-association that serves as an early proxy for high-concentration viscosity and a complementary predictor of in vivo clearance. In a 246-mAb panel, CSI-BLI moderately correlates with viscosity; further, in hFcRn Tg32 mice (41 antibodies), CSI-BLI strongly associates with clearance. Here, we present an end-to-end framework that distinguishes high versus low self-interacting clones (CSI-BLI class) by coupling a fine-tuned protein language model (ESM-2) with residue-aligned 3D context from AlphaFold-predicted structures encoded as residue graphs. Disentangled multi-stream attention fuses sequence content, chain-aware positional information, and structural signals to capture spatially proximate interactions that are distant in sequence. Edit-distance–controlled splits across 1499 IgGs and 988 VHHs assess generalization. The structure-aware model achieves the highest hold-out performance (VHH-Fc *F*1 = 0.76; IgG *F*1 = 0.57), while a sequence-only disentangled variant outperforms a standard PLM baseline without structural inputs. Complementary biophysical feature–based models, built from AlphaFold structures and sequence/structure-derived physicochemical descriptors with cluster-aware selection, deliver robust, interpretable performance (VHH *F*1 = 0.72; IgG *F*1 = 0.57), with SHAP analyses highlighting charge/dipole, hydrophobicity, and aggregation-propensity drivers across CDRs and frameworks. This interaction-aware sequence–structure framework, supported by interpretable feature models, is extensible to other developability endpoints and broader protein classification tasks where joint modeling of language-derived representations and residue-level geometry is advantageous.

## 1 Introduction

Therapeutic antibodies and antibody-derived biologics have transformed modern medicine, but a large fraction of candidate molecules encounter major bottlenecks during development due to unfavorable developability properties such as non-specific binding (polyspecificity), self-association, aggregation, poor solubility/viscosity at high concentration, and manufacturing instability (Jain et al. 2017; Bailly et al. 2020; Zhang et al. 2023). Because these liabilities often emerge after substantial investment in discovery and optimization, there is a strong incentive to identify developability risks as early as possible and to prioritize candidates that are more likely to progress successfully.

CSI-BLI provides a practical, high-throughput measure of antibody self-association using only small amounts of material, making it useful for early screening and candidate down-selection (Sun et al. 2013). In CSI-BLI, antibodies are captured onto AHQ biosensors via the Fc domain, and the remaining capture surface is blocked with free human Fc; self-binding association responses are normalized to an adalimumab negative control to compute fold change, enabling consistent comparative screening in 96-well plates with about 15 *µg* per antibody and 2 hour turnaround. CSI-BLI responses positively correlate with delayed retention times in self-interaction chromatography (SIC) and cross-interaction chromatography (CIC), and with aggregation at high concentration, reinforcing relevance to solubility and viscosity risks driven by weak, reversible self-association (Sun et al. 2013). While AC-SINS and SIC can flag concentration-dependent self-association, they face reproducibility and throughput constraints and can be confounded by column binding artifacts that limit SIC/CIC analyses; in contrast, CSI-BLI directly monitors self-binding on Fc-captured sensors and is plate-based and automation compatible, making it preferable for early discovery screening. Importantly, CSI-BLI is informative beyond self-association per se. Building on the concordance with SIC/CIC and high-concentration aggregation reported by (Sun et al. 2013), we show that CSI-BLI is linked to key downstream liabilities: across a 246-mAb panel, CSI-BLI exhibits a moderate positive association with formulation viscosity and functions as the strongest single-assay classifier of high viscosity under group-aware cross-validation; in pharmacokinetic risk assessment using hFcRn Tg32 mice (Jain et al. 2024), CSI-BLI strongly associates with non-target-mediated clearance. These results position CSI-BLI as a high-throughput anchor for early identification of viscosity and clearance risks, complementary to NSB assays and materially efficient compared with direct high-concentration viscosity measurements or in vivo studies.

However, experimental testing still faces cost, material, and throughput constraints, motivating accurate in silico methods that can pre-screen large libraries, reduce the number of constructs advanced to wet-lab assays, and prioritize engineering efforts. Deep learning has emerged as a promising solution to this problem (Yu et al. 2024; Zhou et al. 2025), but current methods still have limitations. Sequence-based models are fast and scalable. They learn from the amino-acid sequence and can pick up useful signals such as overall composition, recurring motifs, and evolutionary patterns, especially when powered by pretrained protein language models (Rives et al. 2021; Olsen, Moal, and Deane 2022). The downside is that a sequence alone does not explicitly encode 3D geometry, so these models can miss interaction drivers that depend on spatial arrangement, such as surface charge or hydrophobic patches and other structure-dependent effects (Chen et al. 2020; Jain, Boland, and Vásquez 2023; Yu et al. 2024; Éliás et al. 2024). Structure-aware modeling is particularly relevant for CSI-BLI, which detects weak multivalent interactions mediated by spatially clustered surface features.

Structure-based models use 3D information through geometric descriptors or neural networks on molecular surfaces or graphs. This helps them capture spatial interaction cues. But they often need reliable structures, they can be sensitive to how geometry is encoded, and they may not fully use the broad knowledge learned by large sequence models (Rai, Apgar, and Bennett 2023; Park and Izadi 2024; Sormanni et al. 2017). ProSST uses a disentangled attention design to connect sequence tokens with structure-derived tokens during pretraining (Li et al. 2024). While powerful, it operates over a generic sequence–structure token interface. Our approach differs in three concrete ways: (i) we encode structure with a residue-level GVP message-passing encoder on a C_*α*_-based graph, producing per-residue geometric embeddings (scalar and equivariant vector channels) rather than discretized structure tokens; (ii) spatial context is injected through continuous local 3D neighborhoods and relative geometric edge features (e.g., distance RBFs and unit directions), aligned 1:1 with residues; and (iii) we fuse modalities using explicit directional disentangled cross-modal streams (Sequence → Structure and Structure → Sequence) so that sequence representations are updated by residue neighborhoods and, conversely, geometric embeddings are conditioned on sequence context. This design introduces an inductive bias toward spatially clustered surface patterns, which are central to antibody self-association.

We build on CSI-BLI’s practical strengths and the need to reduce wet-lab burden by proposing a residue-aligned, multimodal sequence–structure framework tailored to capture spatial drivers of self-association. Specifically, we couple a fine-tuned protein language model (ESM-2) with AlphaFold-derived geometric embeddings encoded by a GVP graph network and fuse them via disentangled multi-stream attention. By aligning sequence tokens with local geometric neighborhoods, the model targets charge patches and hydrophobic surfaces that can be distant in sequence but contiguous in 3D-patterns directly relevant to CSI-BLI responses.

We propose an end-to-end multimodal model for CSI-BLI class prediction that combines a fine-tuned protein language model with residue-aligned structural context. Instead of simply concatenating sequence and structure features, we use a disentangled attention module that explicitly models how sequence content, positional context, and 3D structures interact. This design lets both sequence and structure representations adapt to the prediction task and provides a more targeted way to capture the interaction patterns that drive antibody self-association.

Complementary to deep learning, biophysical descriptor–based models offer an interpretable pathway to in silico assessment. Using AlphaFold-predicted structures for variable regions and physicochemical descriptors computed with specific software such as Schrödinger, we construct cluster-aware feature sets that mitigate sparsity and multicollinearity and train lightweight classifiers (e.g., SVMs, gradient boosted trees, and soft ensembles). These models achieve robust performance on edit-distance–controlled hold-outs for both VHH and IgG CSI-BLI classification and, via SHAP analysis, reveal mechanistic drivers of self-association-including charge/dipole, hydrophobicity, and aggregation propensity across CDRs and frameworks-providing actionable insights for sequence and surface-property optimization.

Our main contributions are:

- We position CSI-BLI as a high-throughput, low-material anchor for early developability, linking self-association readouts to downstream liabilities: CSI-BLI associates with high-concentration viscosity and provides highest classification performance when combined with non-specific binding assays; similarly, in hFcRn Tg32 mice, CSI-BLI correlates with non-target-mediated clearance and provides complementary information to non-specific binding assays for pharmacokinetic risk.
- We motivate accurate in silico screening to reduce wet-lab burden and propose complementary modeling tracks: (i) interpretable biophysical descriptor models built from AlphaFold structures and sequence/structure-derived physicochemical features with cluster-aware selection, yielding robust performance and SHAP-based mechanistic insights (charge/dipole, hydrophobicity, aggregation propensity across CDRs/frameworks); and (ii) a residue-aligned sequence–structure fusion model that couples fine-tuned PLM embeddings with GVP-encoded geometry.
- We introduce a disentangled multi-stream attention module that fuses sequence content, chain-aware positional information, and structural signals through explicit interaction channels, enabling spatially proximate but sequence-distant features to drive predictions.
- We demonstrate structure-aware gains and generalization across antibody formats using editdistance–controlled splits; the sequence–structure model outperforms sequence-only baselines, and biophysical ensembles offer competitive, robust performance with interpretable drivers.
- We design the architecture in a task-agnostic, modular way (sequence encoder, geometric encoder, disentangled fusion), making it extensible to other developability endpoints and broader protein classification tasks where joint modeling of language-derived representations and residue-level geometry is advantageous.

## 2 Materials and Methods

### 2.1 Experiments and Data Collection

#### 2.1.1 Clone self-interaction by biolayer interferometry (CSI-BLI) measurements

CSI-BLI measurements were performed on an Octet RH16 (Sartorius) with Octet Anti-Human IgG Quantitation (AHQ) Biosensors (Sartorius, Cat# 18-5001). Adalimumab and bococizumab formatted as hIgG1 were used as negative and positive controls, respectively. Test articles and controls were diluted in 1x Octet Kinetics buffer (Sartorius, Cat# 18-1105) to 3 *µ* g/mL for loading and 150 *µg/mL* for association steps. Human Fc fragment (Jackson Immuno Research, 009-000-008) was diluted to 42.4 *µg/mL* in 1x Octet Kinetics buffer for blocking. Sensors were regenerated and neutralized using glycine pH 1.5 (Cytiva, Cat# BR100354) and 1x Octet Kinetics buffer, respectively. Samples were prepared in 384-well tilted bottom Octet plates (Sartorius, Cat# 18-5166) with 50 *µL* per well. Octet AHQ Biosensors were soaked in 1x Octet Kinetics buffer for at least 10 min before measurements. The experiment included baseline (60 sec in 1x Octet Kinetics buffer), loading (3 *µg/mL* antibodies for 100 sec), blocking (human Fc for 180 sec), baseline (60 sec in 1x Octet Kinetics buffer), association (150 *µg/mL* antibodies for 300 sec), dissociation (30 sec in 1x Octet Kinetics buffer), and regeneration. Data was processed in Octet Analysis Studio software with alignment to baseline and Savitzky-Golay filtering applied. CSI-BLI fold change was determined by normalizing the association response to that of the negative control, adalimumab.

#### 2.1.2 Non-specific binding ELISAs

Maxisorp ELISA plates were coated with non-specific binding (NSB) antigens overnight at 4C: BV particles (1%), ssDNA (1 ug/mL Sigma-Aldrich, # D8899-5MG), and cardiolipin (50 ug/mL Sigma-Aldrich # C0563-10MG). Heparin coated plates were used directly from Thermofisher (Part # 50197531). All following steps were performed at room temperature. Wells were washed with PBS and then incubated with blocking buffer (PBS with 0.5% BSA) for 1h, followed by three washes with PBS. Test antibodies diluted in blocking buffer to 15 ug/mL were added to the wells in quadruplicate, incubated for 1 h, then washed 3x with PBS. Goat anti-human IgG-horseradish peroxidase secondary antibodies (1:5000 dilution, Sigma-Aldrich # A0170) in blocking buffer were added to each well and incubated for 1 hr, followed by three washes with PBS. Finally, 3,3’,5,5’-tetramethylbenzidine substrate (SureBlue Reserve, KPL #5120–0081) was added to each well and incubated for 3 min. The reactions were stopped by adding 50of 0.5M sulfuric acid to each well. The absorbance was read at 450nm on a 96-well plate reader (Envision). The assay scores were determined by normalizing absorbance to control wells with no test antibody.

#### 2.1.3 AC-SINS

Briefly, both whole goat IgG (Jackson ImmunoResearch #005-000-003) (non-capture) and polyclonal goat anti-human IgG Fc (Jackson ImmunoResearch #109-005-098) (capture) antibodies were transferred into 20mM potassium acetate (pH 4.3) buffer, and then conjugated to 20nm gold nanoparticles (Innova Biosciences #3201–0100) at a 3:2 ratio of capture:non-capture antibodies. Antibodies were incubated with gold nanoparticles at a 9:1 ratio for 1h at room temperature, and then blocked by the addition of 0.1*µ* M poly-ethylene glycol methyl ether thiol (2000MW, Sigma-Aldrich #729140) for 1h. The coated and blocked nanoparticles were concentrated by centrifugation to an OD535 of 0.4 and stored at 4°C. To assess self-association, 5*µ* L of nanoparticles were mixed with 45*µ* L of purified antibody at 50 ug/mL in PBS, pH 7.2 or HA, (20mM histidine, 200mM arginine) pH 6 in a 384-well plate. Nanoparticles were mixed with buffer only (no antibody) as a control. Absorbance was measured on a SPECTROstar Nano UV/vis plate reader from 490 to 700nm. The wavelength of peak absorbance was calculated in the MARS data analysis software and used to determine the wavelength shift compared to the nanoparticle-only control.

#### 2.1.4 Viscosity Assay

Viscosity of the mAb solutions was measured using MCR203 cone and plate rheometer (Anton Paar, Graz, Austria). Viscosity of each sample (150 mg/mL) was obtained using CP20–1 cone with 19.972 mm diameter that holds 40 *µL* of the sample at a shear rate of 1000 Hz.

### 2.2 CSI-BLI as a High-Throughput Predictor of Viscosity and Clearance

#### 2.2.1 Correlation Analysis

Relationships between assay signals and high concentration formulation viscosity across a 246-mAb panel were quantified using Spearman rank correlation. For each antibody, viscosity values were paired with results from NSB ELISAs (baculovirus particle, cardiolipin, and single-stranded DNA) as well as AC-SINS and CSI-BLI responses. Spearman correlation coefficients were computed for all assay–viscosity pairs in the data set.

#### 2.2.2 Cross validation scheme

Models were evaluated using a leave-one-group-out cross validation approach, to prevent information leakage from closely related antibody variants within the data set. Sequences were partitioned into groups by parental origin. In each fold, one parental group was held out as the test set, with remaining groups acting as the training set. Classifiers were fit on the training data and evaluated on the held out group. Model accuracy, F1, and sensitivity were averaged across folds to estimate performance.

#### 2.2.3 Feature and model selection

Classifier selection was performed across a consistent feature set which included all NSB data, CSI-BLI, and AC-SINS. Maximization of F1 score with minimal model complexity was prioritized during selection, due to limited data set. Multiple models were compared, including linear discriminant analysis (LDA), gradient boosting classifiers, and random forest classifiers. LDA was chosen based on its simplicity, interpretability, and performance comparable to the more complex alternatives.

Following model selection, a full factorial feature screen was conducted to identify the optimal combination of assay inputs for viscosity prediction. The systematic approach was made feasible by the small number of candidate features and samples. Optimal feature set was identified based on maximum cross validated F1.

### 2.3 Machine learning prediction of CSI-BLI

#### 2.3.1 Data distribution, data splitting

##### IgGs

The dataset consists of 1499 sequences from multiple projects, with 30% of sequences belonging to class I, see Fig. 5**a**. Sequence similarity has been shown to substantially impact machine learning model performance in the biologics space. Hence, to prevent data leakage and to ensure proper evaluation of model performance and generalizability, we construct a hold-out test set of 299 diverse sequences that maintain a minimal distance of 20 edit operations from the training set. Specifically, we compute the pairwise Levenshtein distance (Levenshtein 1966) between sequences and use it as the distance metric for an agglomerative clustering algorithm. Using a threshold of 20 edits with single-linkage clustering, we identify “wild-type” sequences in our dataset that are more than 20 edits away from all other sequences, and select a subset of these as the hold-out test set. This selection is stratified to match the target distribution in the training and test sets (30% positive class). The resulting distribution of pairwise distances between training and test sequences is shown in Fig. 5**b**.

##### VHH

The VHH dataset consists of 988 sequences of which 33% belong to Class I (Fig. 5**c**). Analougously to the IgG dataset, we select an hold-out test set of 198 sequences at least 10 edit distance away from all other sequences, with the same approach illustrated for IgG sequences. The resulting distribution of pairwise distance between training and test sequences is shown in Fig. 5**d**.

#### 2.3.2 Biophysical features model

Structure-derived features for each molecule are generated automatically by AstraZeneca’s internal in silico developability pipeline. Sequences are ingested and the variable regions (Fv or VHH) are modeled using AlphaFold2 (Jumper et al. 2021). The structure then serves as the input to multiple tools—MOE 2024:06, Schrödinger 2024-4-js-aws BioLuminate, and CamSol 2.1-foss-2019a-Python-3.7.2—to compute structural descriptors. In parallel, sequence-level descriptors are produced, including sequence liabilities and motifs of interest. These features will be used for model training.

#### 2.3.3 Cross validation scheme

To choose the best model configurations and parameters, we adopt a cross-validation scheme based on the clusters obtained as explained in section 2.3.1. Specifically, we use the 5-StratifiedGroupKFold methods from Scikit-Learn using the sequence clusters obtained with hierarchical clustering as grouping parameter. This ensures that clusters appear either in train or test in the different folds, ensuring that no data leakage occurs. To select the best model configuration, we concatenate the out-of-fold predictions and select the configurations with highest F1-score on the validation set.

#### 2.3.4 Feature and model selection

As the number of available physicochemical descriptors is large (*>* 1000 available features), we implement a feature selection strategy to increase the robustness of the model. To increase generalizability, we remove features from granular regions of the antibodies, such as specific CDR/Framework subregions. To mitigate multicollinearity, we remove features with a Spearman correlation larger than 0.8. Furthermore, we exclude features with more than 80% null values and those with a coefficient of variation smaller than 0.1. This results in an initial pool of features of 164 and 93 descriptors for IgGs and VHHs, respectively. The subset is further refined using a univariate, cluster-aware, validated selection. Specifically, we create *M* folds iteratively holding out each of the *M* clusters identified by the method described in Section 2.3.1. For each fold, we obtain the set of *k* features with the largest ANOVA F-scores with respect to the target. To establish the final subset, we select the *k* features that appear most frequently across folds. For each candidate model, we then choose *k* as the feature set that achieves the highest F1-score using the cross-validation scheme described in Section 2.3.3.

#### 2.3.5 Hyperparameter tuning

Once the optimal number of features for each candidate model is selected following the procedure described in the previous section, we perform hyperparameter tuning with grid search. Parameters explored for each model are listed in 6. The best model configuration is chosen according to the highest F1-score on concatenated validation predictions.

#### 2.3.6 Shap values and feature importance

To assess potential mechanisms underlying self-association risk, we use the widely adopted SHAP library for feature importance (Lundberg et al. 2020). Specifically, we use the SHAP package to compute the feature importance of the Gradient Boosted Tree model within the Ensemble model on the training data. To identify clusters of Class I sequences, we apply the HDBSCAN clustering algorithm (Campello, Moulavi, and Sander 2013) to the space defined by the SHAP values of Class I sequences, akin to a *supervised*-clustering approach. In this analysis, we define *positive clusters* as those comprising sequences annotated as Class I, whereas the *negative cluster* corresponds to Cluster 0. Sequences labeled as noise by the algorithm are reassigned in a subsequent step to the closest cluster in shap space. For HDBSCAN hyperparameter selection, we track multiple clustering metrics (Silhouette score, Davies–Bouldin index, and Calinski–Harabasz index) and choose the settings that optimise these metrics (see Fig. 7). To quantify feature differences, we test for features that are significantly different in each *positive cluster* relative to the *negative cluster* using the Kruskal–Dunn test with Bonferroni correction for multiple comparisons. We then select the top-3 most significantly different features (lowest adjusted p-value) for each cluster, which results in total to 9 unique significantly different features (shown in supplementary Fig. 9). Four features with the largest mean shap value are then shown in Fig. 3.

### 2.4 PLM/GNN model architecture

Our goal is to predict clone self-interaction from paired heavy/light chain (IgGs) or single-chain VHH sequences and their corresponding three-dimensional structures. To this end, we develop a joint protein language model (PLM) and geometric vector perceptron (GVP) architecture with a disentangled multi–stream attention mechanism. The model operates at the residue level and aggregates sequence, structural, and positional information into a single antibody-level representation used for binary classification (Figure 1).

**Figure 1:**
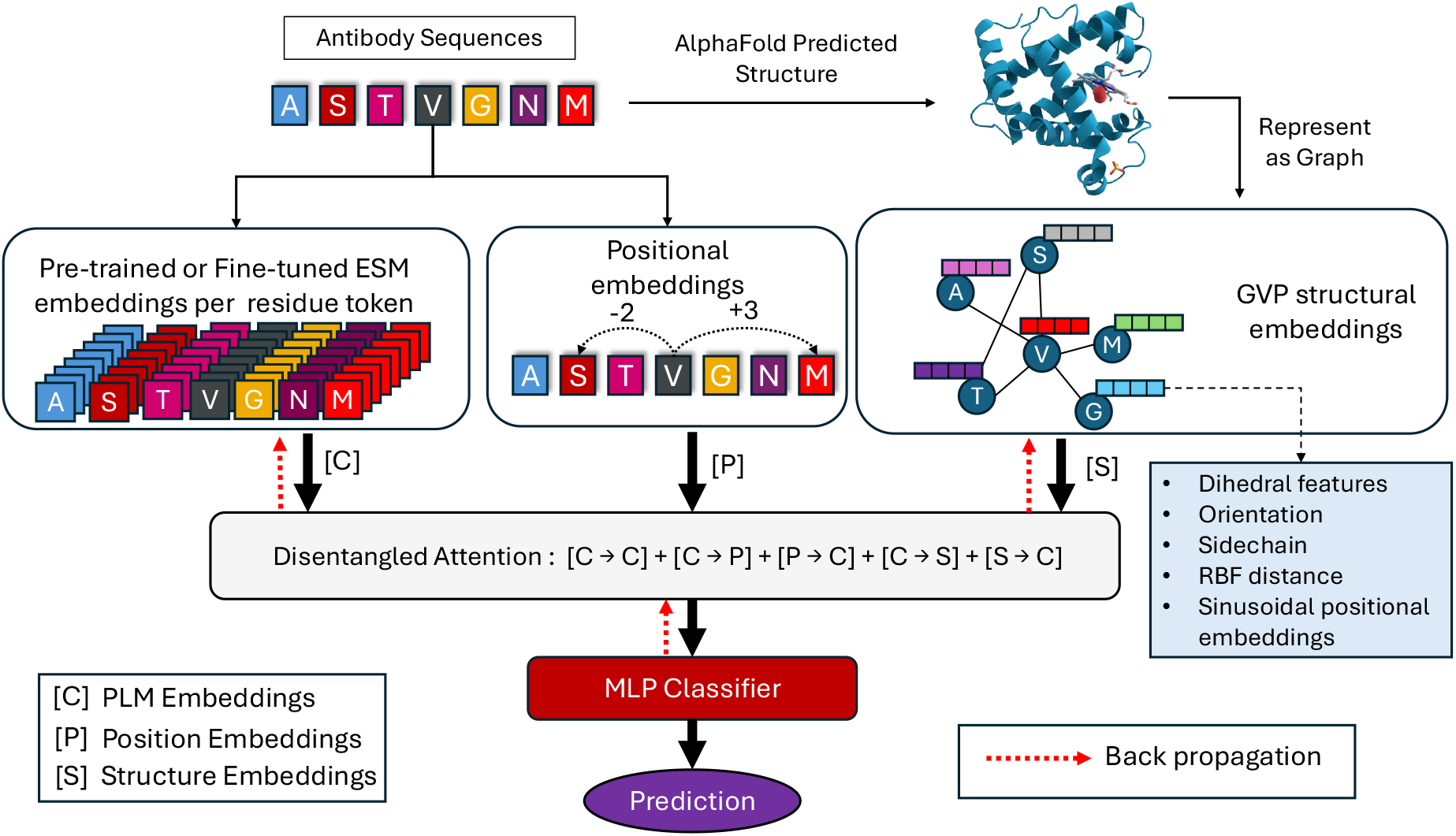
Overview of the disentangled PLM–GNN architecture for antibody residue-level representation learning. Pre-trained or fine-tuned ESM models generate per-residue content embeddings [C], while positional embeddings [P] encode relative sequence context. Structural information [S] is derived from a Geometric Vector Perceptron (GVP) encoder operating on residue graphs with di-hedral angles, backbone orientation, side-chain direction, RBF-expanded distances, and sinusoidal positional encodings. The three streams - content, position, and structure are fused through a disentangled attention mechanism that separately models interactions such as C→C, C→P, P→C, C→S, and S→C. The resulting multimodal representation is aggregated and passed to an MLP classifier to produce the final prediction. Backpropagation updates both the PLM (in full fine-tuning or LoRA mode) and the GVP encoder, enabling end-to-end learning.

#### 2.4.1 Protein Language Model (PLM)

We use an ESM-2 encoder ℰ_PLM_ (`facebook/esm2_t33_650M_UR50D`) as the sequence encoder. For IgG antibodies, we encode the heavy and light chain amino acid sequences (*s*^VH^, *s*^VL^) separately:

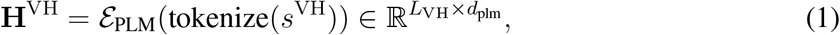

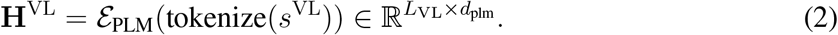

We remove special tokens (e.g., CLS/EOS) by discarding the first and last positions of the PLM output, and concatenate residue embeddings from the two chains along the sequence dimension:

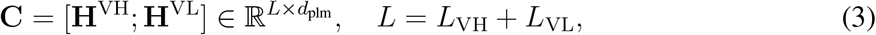

which we refer to as the *content* stream. For VHH antibodies (single-domain heavy-chain-only), there is no light chain sequence; we therefore set **C** = **H**^VH^ (i.e., we bypass the concatenation step) and apply the same downstream architecture unchanged.

#### 2.4.2 PLM Fine-tuning Strategies

We evaluate three PLM fine-tuning modes: **Frozen** (PLM weights fixed; only downstream modules are trained), **Full** (all PLM parameters are updated end-to-end), and **LoRA** (train low-rank adapter parameters on selected attention projections while keeping the base PLM frozen).

#### 2.4.3 Structure Graph Construction

For each antibody, we have the corresponding AlphaFold predicted PDB structure with heavy and light chains. We extract backbone atom co-ordinates **X** ∈ ℝ^*N* ×3×3^ for *N* residues, where the last two dimensions correspond to atoms (N, C_*α*_, C). Non-standard residues are discarded.

We construct a residue level graph *G* = (*V, E*) with |*V* | = *N* nodes. Nodes correspond to residues, and edges (*i, j*) ∈ *E* are built using a *k* nearest neighbor strategy over C_*α*_ co-ordinates:

- For each node *i*, we compute Euclidean distances 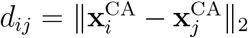 to all other residues.
- We restrict candidates to a radius *d*_*ij*_ ≤ *r* and choose up to *k* nearest neighbors. If fewer than a minimum *k*_min_ neighbors satisfy the radius constraint, we add additional neighbors by nearest distance until *k*_min_ is reached.

##### Node Features

We follow a standard backbone-geometry parameterization and construct per-residue scalar and vector features from the N–C_*α*_–C backbone atoms. Specifically, for each node we compute the backbone dihedrals torsion angles (scalar features), backbone and side chain orientation vectors (vector features). Details on the node features are provided in the supplementary information.

##### Edge Features

For each edge (*i, j*), we compute:

- **Distance RBF features (scalar)**. The C_*α*_ distance 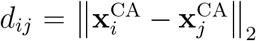 is expanded using radial basis function (RBF) encoding:

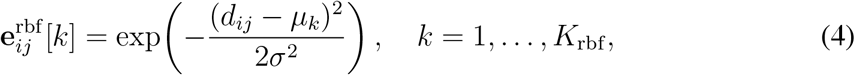

where {*µ*_*k*_} are evenly spaced centers over a fixed interval and *σ* is the bin width.
- **Relative positional encoding (scalar)**. We encode the sequence index difference Δ_*ij*_ = *i* − *j* using sinusoidal embeddings:

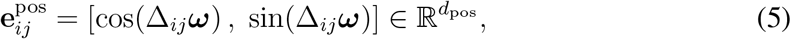

where ***ω*** is a set of frequencies as in Transformer positional encodings.
- **Direction vector (vector)**. We use the unit direction vector from residue *j* to residue *i*,

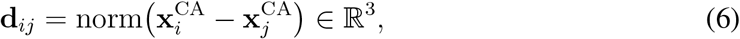

stored as a single-channel vector edge feature.

We concatenate scalar edge features

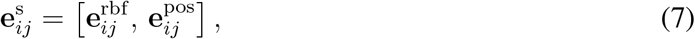

and set the vector edge feature to

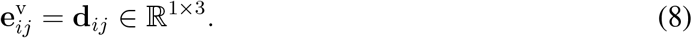

#### 2.4.4 Geometric Vector Perceptron (GVP)

We use a GVP message passing network (Jing et al. 2020) to jointly process scalar and vector features associated with each residue. Each node *i* is represented by scalar features 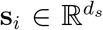 and vector features 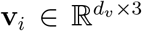. A GVP module applies separate transformations to the two feature types,

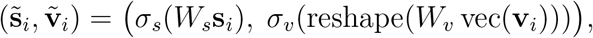

ensuring that vector information is updated in a rotation-aware manner. For each edge *i* → *j*, node and edge features are processed through dedicated GVPs and combined into a single message,

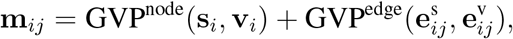

which is then aggregated over neighbors by summation:

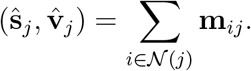

Residual connections are used to stabilize training and preserve information across layers. Stacking *L*_GVP_ layers gives refined per residue representations 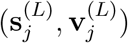. We convert them to a unified scalar structural embedding by concatenating scalar outputs with vector norms,

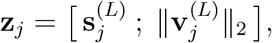

resulting in the structural stream 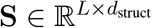 used by the multimodal attention module.

#### 2.4.5 Positional Information

To encode residue order within our disentangled attention, we evaluate two strategies: (i) an *absolute positional stream* that explicitly represents residue indices and participates in cross-stream interactions, and (ii) a *relative positional formulation* that injects pairwise offset information directly into the attention logits (DeBERTa-style). For IgGs, the heavy (VH) and light (VL) chains are distinct physical entities; although we concatenate their residue embeddings into a single tensor for implementation convenience, positional information is computed in a *chain-aware* manner so that the VH–VL boundary is not treated as adjacency.

##### Chain-aware indexing

Let **C** = [**H**^VH^; **H**^VL^] ∈ ℝ^*L*×*d*^ with *L* = *L*_VH_+*L*_VL_. For *i* ∈ {0, …, *L*− 1},

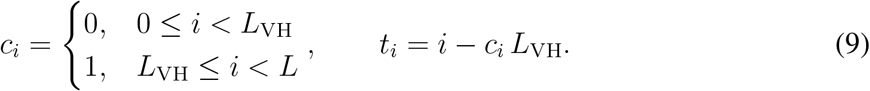

Thus, both chains start at *t* = 0 in their own coordinate system.

##### Absolute positional stream

Given an embedding table ℰ_pos_ shared across samples, we embed positions using the within-chain index and add a learned chain embedding:

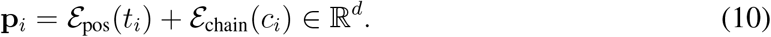

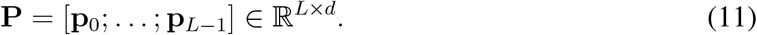

This produces an explicit positional representation aligned with the content and structure streams and enables CP and PC interaction channels within the disentangled attention module, without imposing an artificial contiguity between the VH and VL termini.

##### Relative positional offsets (DeBERTa-style)

Alternatively, we encode position through relative displacements computed from within-chain indices. For residues *i* and *j*, we define

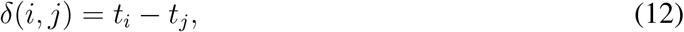

clip to a bounded range [−(*M* − 1), …, (*M* − 1)], and look up a learned relative embedding:

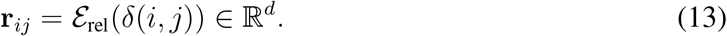

To explicitly distinguish inter-chain from intra-chain pairs, we also add a learned chain-pair embedding (or bias) based on (*c*_*i*_, *c*_*j*_):

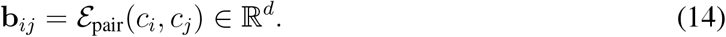

The relative term is then 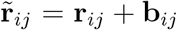. The same base embedding is projected into *key-role* and *query-role* components,

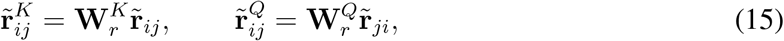

and contributes additively to the attention logits via content-relative and relative-content terms. This removes the need to construct an explicit **P** stream and provides translation-invariant positional information *within* each chain while preserving a distinct treatment of VH–VL interactions.

#### 2.4.6 Disentangled Multi-Modal Attention

Let **C** ∈ ℝ^*L*×*d*^ denote content embeddings (sequence-derived token features) and **S** ∈ ℝ^*L*×*d*^ denote structure embeddings (residue-aligned 3D features). Depending on the positional strategy above, we either use an absolute positional stream **P** ∈ ℝ^*L*×*d*^ or relative positional embeddings **r**_*ij*_.

For each head *h* with dimension *d*_*h*_, we obtain stream-specific queries/keys by linear projection,

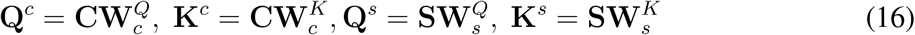

and use content-derived values

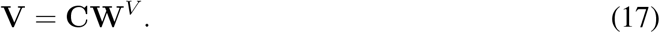

##### Five-channel formulation with absolute positions

When using absolute positions, we additionally project **P** into (**Q**^*p*^, **K**^*p*^). The attention logit between residues *i* and *j* is decomposed into five interaction channels:

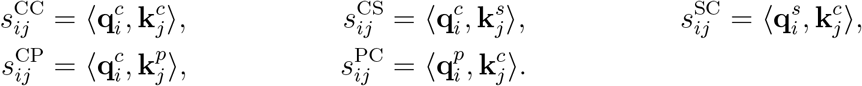

The combined score is

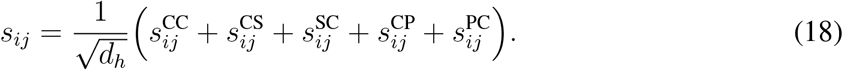

##### Relative-position formulation within logits

When using relative positions, we omit **P** and instead add DeBERTa-style relative terms to the content-based attention. Specifically,

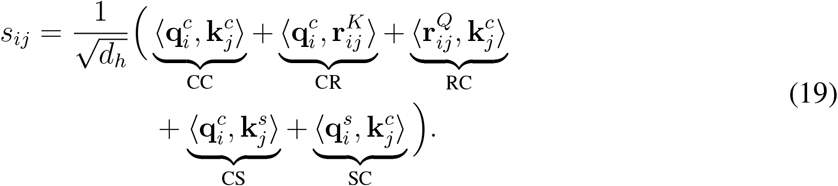

thereby injecting positional information through relative displacements while retaining the same cross-stream content–structure coupling (CS, SC).

##### Attention output and pooling

For both variants, padded residues are masked before softmax:

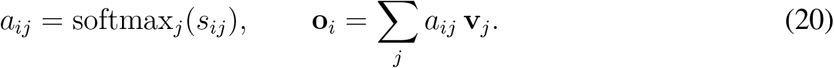

Outputs from all heads are concatenated and projected to produce residue-level fused embeddings **O** ∈ ℝ^*L*×*d*^. We obtain a sequence-level representation via masked mean pooling over valid residues, followed by a final linear layer to produce prediction logits for clone self-interaction inference.

#### 2.4.7 Hyperparameters

Table 1 summarizes the hyperparameters and architectural choices considered in our experiments. We varied how the PLM was adapted (from a frozen backbone to parameter-efficient LoRA updates and full fine-tuning) and compared two objective functions designed for class imbalance. For the structure branch, we evaluated alternative graph neighborhood definitions (top-*k* vs. radius) and varied the depth of the GVP-based encoder. Finally, we tested positional information using either absolute embeddings or DeBERTa-style relative positional encoding to assess how different position parameterizations affect performance.

**Table 1:** Hyperparameters and design choices explored during model development.

| Category | Options |
| --- | --- |
| PLM adaptation strategy | Frozen encoder; LoRA fine-tuning; Full fine-tuning |
| Loss function | Class-weighted cross-entropy; Focal loss |
| Structure graph construction | Top-$k$ nearest neighbors; Distance radius |
| Structure encoder depth | Number of GNN (GVP) layers |
| Positional encoding | Absolute positional embeddings; DeBERTa-style positional encoding |

#### 2.4.8 Computational Resources

We use ESM2 (650M) with a GVP geometric encoder and disentangled multi-stream attention. Training is GPU-intensive; we recommend an NVIDIA A100 (80 GB) or similar for batch size ≥ 16. CPU needs are modest (4–8 cores) for preprocessing/data loading, with peak RAM typically 40–55 GB depending on graph and sequence sizes. Full *K*-fold runs (graph construction, embedding extraction, repeated training) are compute-heavy and can take hours per configuration.

## 3 Results

### 3.1 CSI-BLI as a High-Throughput Predictor of Viscosity and Clearance

#### 3.1.1 Prediction of Viscosity by CSI-BLI Assay

High formulation viscosity at manufacturing concentrations leads to poor syringeability and slow fill–finish operations. Mechanistically, elevated viscosity is driven by reversible self-association, and to a lesser degree, NSB interactions. CSI-BLI response measuring self-association is positively correlated with high-concentration viscosity and aggregation metrics (Kelly et al. 2015), making it a suitable proxy for viscosity risk in early developability, where direct measurement of formulation viscosity is prohibited by material limitations. AC-SINS (affinity-capture self-interaction nanoparticle spectroscopy) has been similarly leveraged as an early-stage proxy for formulation viscosity risk (Armstrong et al. 2024). While informative, AC-SINS can exhibit high variability due to batch-to-batch variation of the gold capture nanoparticles, and has lower throughput with higher material consumption. By contrast, CSI-BLI uses all defined, commercial reagents and requires minimal material, making it more desirable for rapid, scalable screening in early discovery.

While reversible self-association is the primary driver of elevated viscosity at manufacturing concentrations, non-specific interactions with excipients, host cell proteins, and surfaces can further increase viscosity. Early stage screens leverage polyspecificity ELISAs to detect binding propensity to unrelated antigens, including Single-stranded DNA (ssDNA), Cardiolipin, and Baculovirus Particles (BVP). High-throughput adapted ELISA provides a material efficient complement to self-association to characterize formulation viscosity risk in early developability.

Spearman rank analysis on a 246 mAb dataset reveals a moderate positive relationship between viscosity reversible self-association, yielding correlation coefficients of *ρ* = 0.35 and *ρ* = 0.34 for CSI-BLI and AC-SINS, respectively. A higher correlation is observed between CSI-BLI and the non-specific binding data (BVP *ρ* = 0.75; ssDNA *ρ* = 0.60; Cardiolipin *ρ* = 0.60), suggesting that self-association is likely to co-occur with non-specific binding behavior, exacerbating formulation viscosity challenges.

Linear discriminant analysis (LDA) classifiers were trained on assay data to predict antibody viscosity at high-concentration formulations, with leave-one-parental-group-out cross-validation to prevent data leakage among related sequences. Feature combinations of three or more assays were benchmarked against a majority-class null model (Accuracy ≈ 0.72). Prioritizing F1 as the primary ranking metric, the top configurations combined non-specific binding readouts with self-association assays: the highest performing model leveraged Baculovirus Particle (BVP) ELISA, Cardiolipin ELISA, ssDNA ELISA, AC-SINS, and CSI-BLI as features, achieving F1 = 0.57, Accuracy = 0.86, and Sensitivity = 0.52, see table 5.2. Across feature configurations, these results underscore that combining broad non-specific binding readouts with self-association measures yields the most reliable classification of high viscosity risk.

**Table 2:** Hold-out test set performance of biophysical features models for IgG and VHH datasets.

| Data | Model | Acc | Prec | Rec | F1 | ROC | AUPRC | MCC |
| --- | --- | --- | --- | --- | --- | --- | --- | --- |
| **VHH** | Ensemble | **0.8059** | 0.6719 | **0.7818** | **0.7227** | **0.8515** | **0.7247** | **0.5786** |
|  | SVM | 0.7824 | **0.6957** | 0.5818 | 0.6337 | 0.8225 | 0.6824 | 0.4845 |
| **IgG** | Ensemble | **0.6990** | **0.4958** | **0.6629** | **0.5673** | **0.7612** | **0.5649** | **0.3524** |
|  | Gradient Boosted Tree | 0.6321 | 0.4245 | **0.6629** | 0.5175 | 0.7208 | 0.5432 | 0.2585 |

#### 3.1.2 Prediction of Tg32 Clearance by CSI-BLI

Clearance (CL) arises from antibody interactions with the endothelial glycocalyx and cellular uptake pathways coupled to FcRn-mediated recycling, and is distinct from target-mediated drug disposition (Bryniarski et al. 2026; Datta-Mannan et al. 2015). Elevated non-target-mediated CL reduces systemic exposure and half-life, complicates dose selection, narrows the therapeutic window, and increases manufacturing burden. From a developability perspective, early identification of molecules predisposed to fast CL enables sequence or surface property optimization prior to resource-intensive in vivo studies.

Here, we leveraged a published dataset comprising 43 human IgG1 antibodies raised against hen egg lysozyme (Jain et al. 2024). The panel was selected to span diverse human germlines and physicochemical properties while avoiding mouse cross-reactivity. Linear clearance (dose-normalized AUC) was measured following a single intravenous dose (5 mg/kg) in hFcRn Tg32 homozygous mice (Jain et al. 2024). In a pilot replication focused on high-throughput assays, we reproduced the 43-antibody panel and evaluated four assays: nonspecific binding ELISAs (BVP variants, ssDNA and Cardiolopin) and CSI-BLI. To ensure sequence diversity, we assessed pair-wise differences and found that most molecules differed by more than 15 amino acid substitutions, while two pairs were separated by only three mutations. One member of each closely related pair (ADI-66788, ADI-66789) was excluded, resulting in a final dataset of 41 antibodies.

In our analysis, CSI-BLI shows a strong association with Tg32 linear clearance (Spearman *ρ* ≈ 0.65; p *<<* 0.01), placing it among the top single-assay predictors reported in (Jain et al. 2024)’s paper. Its performance is comparable to the leading nonspecificity assays PSR (*ρ* = 0.69) and BVP (*ρ* = 0.67) and clearly outperforms the self-association assay AC-SINS (*ρ* = 0.55) and classical developability metrics such as HIC (*ρ* = 0.39).

### 3.2 Machine learning prediction of CSI BLI

### 3.3 Biophysical feature-based models

We evaluate biophysical feature–based models for VHH and IgG CSI-BLI classification. Following the validation procedure described in Sec.2.3, we benchmark a set of candidate classifiers on the validation set and report the best-performing configurations in Table 7. Overall, the VHH dataset appears more separable, with consistently higher performance across metrics, indicating that the selected physicochemical descriptors capture a clearer signal for VHH self-association. In contrast, IgG classification is more challenging and exhibits a less favorable precision–recall balance, consistent with greater sequence/format complexity and potentially more heterogeneous mechanisms contributing to self-interaction in full-length antibodies.

Across both datasets, SVM and Gradient Boosted Trees (GBT) achieve the highest scores on selected metrics (e.g., Accuracy/MCC for SVM and AUPRC for GBT in VHH), highlighting their predictive strength; however, no single model dominates across all metrics. To prioritize robustness, we also evaluate an Ensemble that combines the outputs of all single models via soft averaging. The Ensemble delivers consistently strong, balanced performance and never yields the lowest value on any metric, indicating robust generalization across tasks and evaluation criteria for both IgG and VHH. Based on these observations, for each modality we select the top-performing model by F1-score alongside the Ensemble to assess generalization performance on the hold-out unseen test set.

The performance metrics of the top-performing models on the hold-out test set are reported in Table 2. In both modalities, the Ensemble consistently improves aggregate measures of classification quality—particularly F1, MCC, and AUPRC—relative to the single best model (e.g., VHH F1: 0.7227 vs. 0.6337; MCC: 0.5786 vs. 0.4845), indicating that averaging the strengths of individual predictors yields a more robust and generalizable model. Both models are capable of ranking sequences, as shown by ROC–AUC substantially above chance and corroborated by AUPRC (see Fig. 2**b**,**d**).

**Figure 2:**
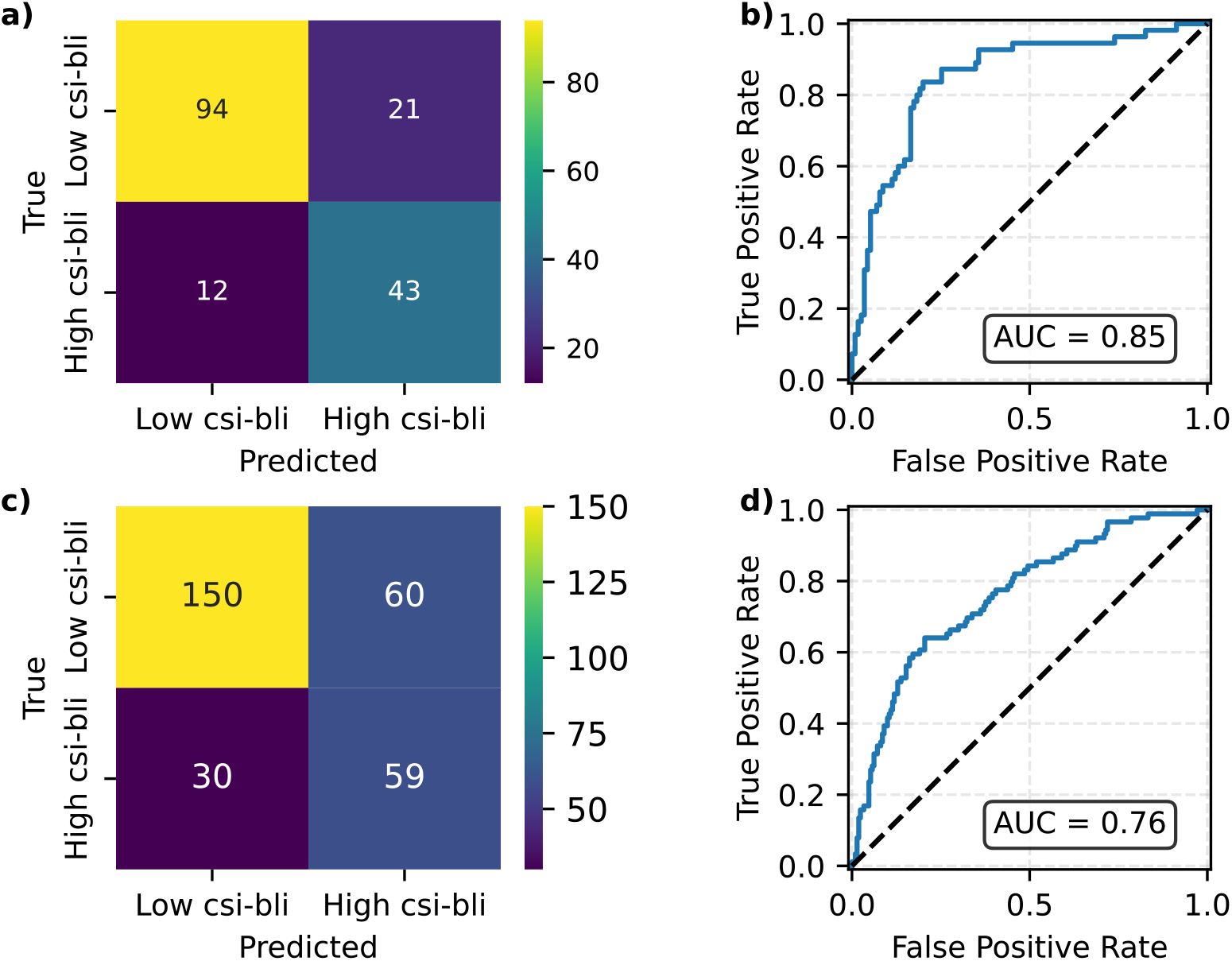
Confusion matrix and ROC curve on the VHH (**a**,**b**) and IgG (**c**,**d**) test sets for the Ensemble biophysical features models.

**Figure 3:**
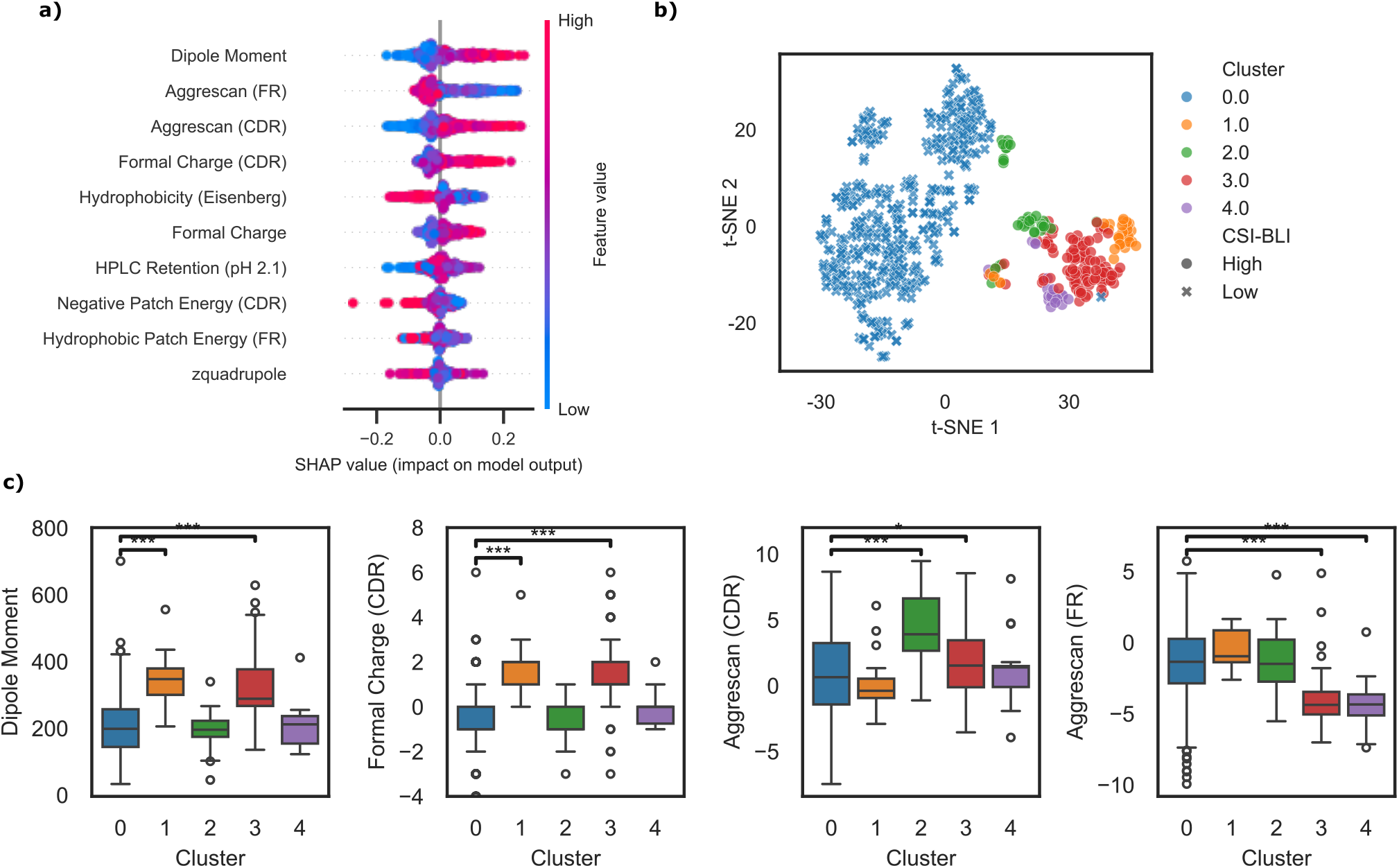
Shap values reveal different biophysical drivers of high CSI-BLI in VHHs.**a**, Shap values for the 10 most important features on the training set. **b)**, training sequences on a 2D t-SNE projection of SHAP explanation space. Colors indicate clusters of sequences with different biophysical CSI-BLI drivers, while the symbol wether the sequences has high or low CSI-BLI. **c**, Distribution of four selected features for the different clusters found in the SHAP (explanation) space. Significance shown only on comparison between clusters 1-4 (high CSI-BLI) and cluster 0 (low CSI-BLI). Stars indicate significance according to Kruskal-Dunn test with Bonferroni correction for multiple comparisons. ∗∗∗ : *p <* 0.001.

### 3.4 Transformer-based models

Tables 3 and 8 summarize hold-out test performance and cross-validation results for IgG and VHH across the three model variants.

**Table 3:**
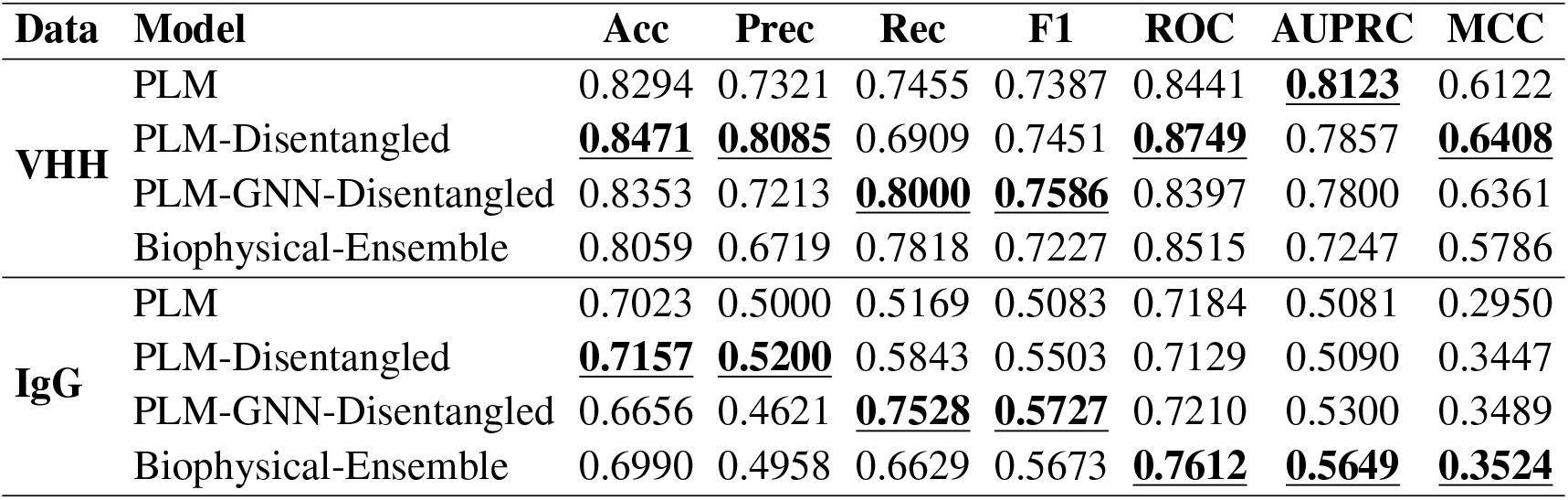
Hold-out test set performance of the models for IgG and VHH datasets for transformer-based models. Performance of the ensemble models from the previous section is added to the table for completeness.

| Data | Model | Acc | Prec | Rec | F1 | ROC | AUPRC | MCC |
| --- | --- | --- | --- | --- | --- | --- | --- | --- |
| VHH | PLM | 0.8294 | 0.7321 | 0.7455 | 0.7387 | 0.8441 | **0.8123** | 0.6122 |
|  | PLM-Disentangled | **0.8471** | **0.8085** | 0.6909 | 0.7451 | **0.8749** | 0.7857 | **0.6408** |
|  | PLM-GNN-Disentangled | 0.8353 | 0.7213 | **0.8000** | **0.7586** | 0.8397 | 0.7800 | 0.6361 |
|  | Biophysical-Ensemble | 0.8059 | 0.6719 | 0.7818 | 0.7227 | 0.8515 | 0.7247 | 0.5786 |
| IgG | PLM | 0.7023 | 0.5000 | 0.5169 | 0.5083 | 0.7184 | 0.5081 | 0.2950 |
|  | PLM-Disentangled | **0.7157** | **0.5200** | 0.5843 | 0.5503 | 0.7129 | 0.5090 | 0.3447 |
|  | PLM-GNN-Disentangled | 0.6656 | 0.4621 | **0.7528** | **0.5727** | 0.7210 | 0.5300 | 0.3489 |
|  | Biophysical-Ensemble | 0.6990 | 0.4958 | 0.6629 | 0.5673 | **0.7612** | **0.5649** | **0.3524** |

**Table 4:** Top 10 LDA models ranked by F1 for predicting high-concentration viscosity using leave-one-parental-group-out cross-validation.

| Assays | Accuracy | F1 Score | Sensitivity |
| --- | --- | --- | --- |
| BVP ELISA, Cardiolipin ELISA, CSI-BLI, AC-SINS | 0.8546 | **0.5692** | 0.5290 |
| BVP ELISA, Cardiolipin ELISA, ssDNA ELISA, CSI-BLI, AC-SINS | 0.8546 | **0.5692** | 0.5290 |
| BVP ELISA, Cardiolipin ELISA, ssDNA ELISA, AC-SINS | 0.8561 | 0.5687 | 0.5224 |
| BVP ELISA, Cardiolipin ELISA, CSI-BLI | 0.8517 | 0.5636 | 0.5328 |
| BVP ELISA, ssDNA ELISA, AC-SINS | 0.8485 | 0.5535 | 0.5224 |
| BVP ELISA, Cardiolipin ELISA, AC-SINS | 0.8449 | 0.5502 | 0.5224 |
| BVP ELISA, Cardiolipin ELISA, ssDNA ELISA, CSI-BLI | 0.8412 | 0.5467 | 0.5328 |
| BVP ELISA, Cardiolipin ELISA, ssDNA ELISA | 0.8381 | 0.5278 | 0.5036 |
| BVP ELISA, ssDNA ELISA, CSI-BLI | 0.8493 | 0.5277 | 0.4733 |
| BVP ELISA, ssDNA ELISA, CSI-BLI, AC-SINS | 0.8450 | 0.5200 | 0.4672 |

**Table 5:** Hyperparameters explored for each model configuration for VHH. In bold the final selected hyperparameters.The Naive Bayes model (not listed here) used 10 features.

| Model | Features | Hyperparameters explored |
| --- | --- | --- |
| **Logistic Regression** | 35 | penalty = **elasticnet**; C = [0.01, **0.1**, 1.0, 10]; solver = **saga**; max_iter = **2000**; l1_ratio = [**0.1**, 0.2, 0.3, 0.4, 0.5, 0.6, 0.7, 0.8, 0.9, 1.0] |
| **SVM** | 20 | C = [0.01, 0.1, **1.0**, 10]; kernel = ["linear", "**rbf**", "poly"]; gamma = ["**scale**", "auto"] |
| **Gradient Boosted Tree** | 20 | max_iter = [**100**, 200]; learning_rate = [0.01, **0.1**]; max_depth = [3, **5**, 7]; min_samples_leaf = [**10**, 20, 30]; max_features = [**0.9**, 1.0] |

**Table 6:**
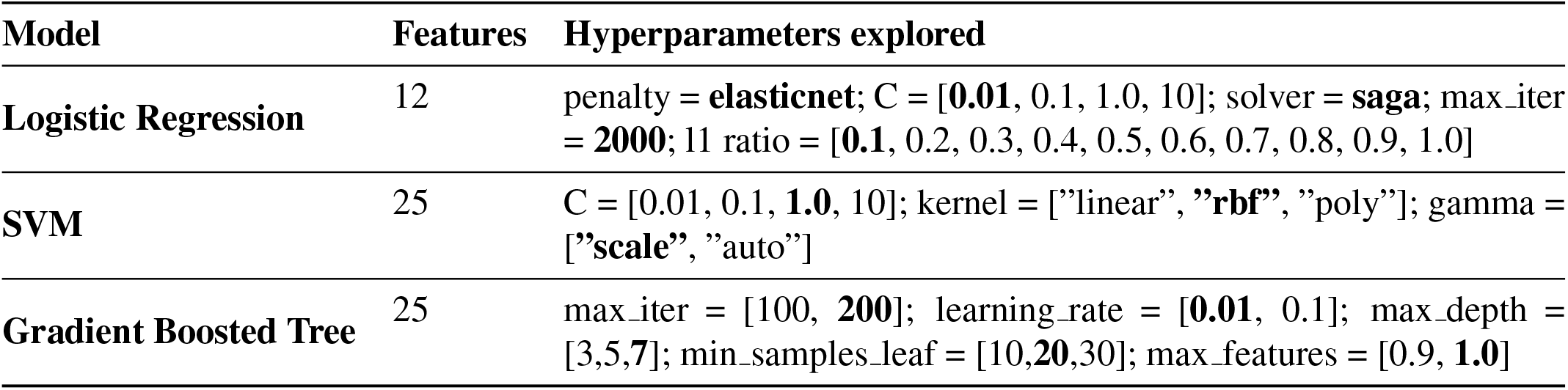
Hyperparameters explored for each model configuration for IgG. In bold the final selected hyperparameters. The Naive Bayes model (not listed here) used 20 features.

**Table 7:**
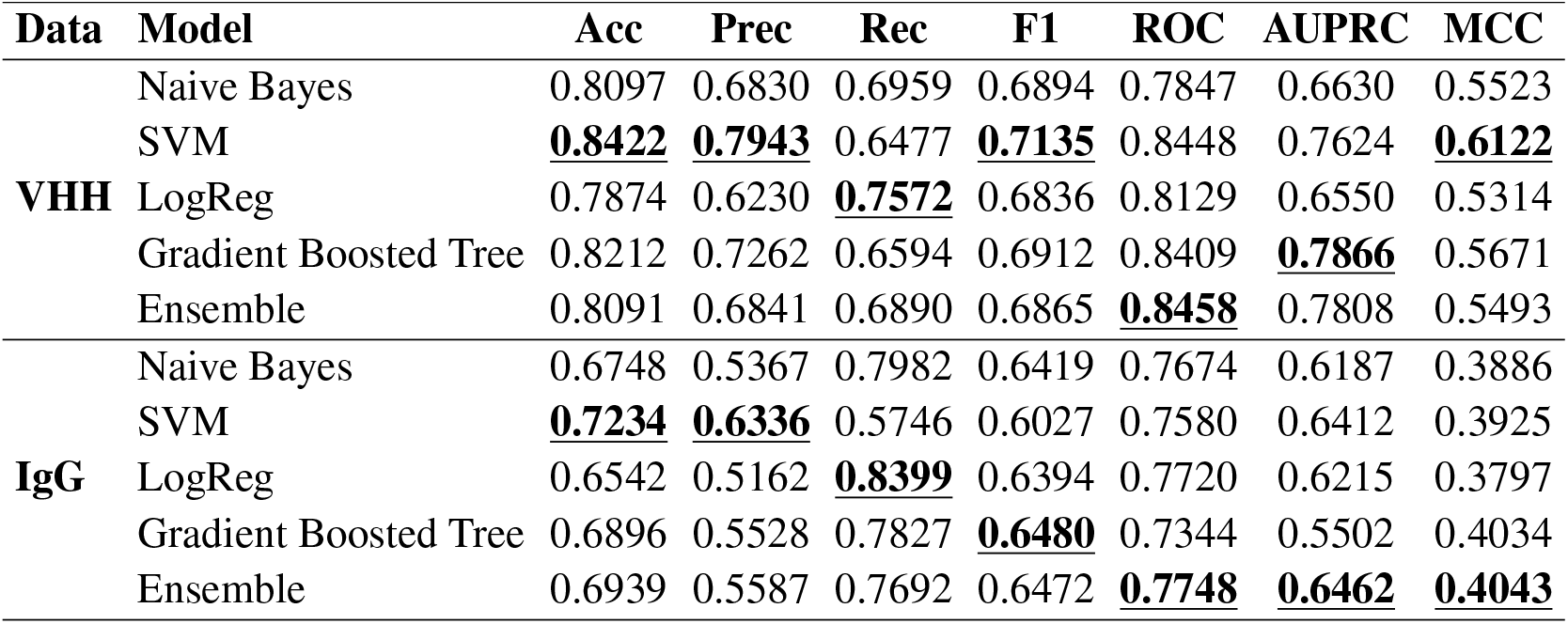
Validation performance of biophysical features models for IgG and VHH datasets.

**Table 8:** Cross-validation performance (mean ± std) of the models for IgG and VHH datasets. Results are averaged across predefined folds and reported as mean ± standard deviation.

| Data | Model | Acc | Prec | Rec | F1 | ROC | AUPRC | MCC |
| --- | --- | --- | --- | --- | --- | --- | --- | --- |
| VHH | PLM | <u>.86</u> ± 0.03 | <u>.83</u> ± 0.15 | .66 ± 0.09 | .73 ± 0.09 | <u>.87</u> ± 0.06 | <u>.81</u> ± 0.14 | <u>.62</u> ± 0.07 |
| | PLM-Disentangled | .83 ± 0.07 | .69 ± 0.10 | <u>.82</u> ± 0.15 | <u>.74</u> ± 0.10 | .83 ± 0.10 | .73 ± 0.19 | .50 ± 0.27 |
| | PLM-GNN-Disentangled | .83 ± 0.04 | .71 ± 0.17 | .71 ± 0.12 | .71 ± 0.13 | .82 ± 0.06 | .72 ± 0.16 | .56 ± 0.11 |
| IgG | PLM | .66 ± 0.06 | .50 ± 0.08 | .82 ± 0.08 | .62 ± 0.05 | <u>.76</u> ± 0.02 | .59 ± 0.04 | .39 ± 0.04 |
| | PLM-Disentangled | .67 ± 0.09 | .51 ± 0.04 | <u>0.86</u> ± 0.09 | .64 ± 0.03 | .75 ± 0.08 | .56 ± 0.07 | .40 ± 0.10 |
| | PLM-GNN-Disentangled | <u>.68</u> ± 0.11 | <u>.54</u> ± 0.08 | 0.82 ± 0.09 | <u>.64</u> ± 0.05 | 0.76 ± 0.07 | <u>.60</u> ± 0.11 | <u>.41</u> ± 0.13 |

On the VHH hold-out test set (Table 3), PLM-GNN-Disentangled attains the highest F1 (0.7586) and recall (0.8000), while the PLM baseline achieves the best AUPRC (0.8123). PLM-Disentangled yields the best accuracy (0.8471), precision (0.8085), ROC-AUC (0.8749), and MCC (0.6408). Overall, these results suggest complementary strengths: structure-aware modeling improves sensitivity and F1 on this split, whereas disentangled fusion improves discrimination across several threshold-dependent metrics, and the PLM baseline remains highly competitive in ranking quality under class imbalance. The best-performing PLM-GNN-Disentangled configuration used a frozen PLM with a StructGVP module, cross-entropy loss, batch size 16, learning rate 1 × 10^−4^, and 20 epochs. For graph construction, we used a kNN radius of 20 Å, selected at most 10 neighbors per node (top-k = 10), and enforced a minimum of 1 neighbor per node (min-k = 1). The model used 3 hidden layers with hidden and embedding dimensions of 256, and dropout 0.1. On VHH data, PLM-based models match or outperform the biophysical feature–based ensemble, with higher F1 and Recall for PLM-GNN-Disentangled and higher ROC for PLM-Disentangled.

On the IgG hold-out test set (Table 3), PLM-GNN-Disentangled achieves the best F1 (0.5727), recall (0.7528), ROC-AUC (0.7210), AUPRC (0.5300), and MCC (0.3489), while PLM-Disentangled improves over the PLM baseline in F1 (0.5503 vs. 0.5083) and MCC (0.3447 vs. 0.2950). This pattern indicates that disentangled attention provides consistent gains over the sequence-only baseline, and incorporating residue-level geometric context further improves overall performance on the IgG test split. The best-performing PLM-GNN-Disentangled configuration used LoRA-enabled PLM fine-tuning with a StructGVP module, focal loss, batch size 16, learning rate 1 × 10^−4^, and 20 epochs. For graph construction, we used a kNN radius of 20 Å, selected at most 30 neighbors per node (top-k = 30), and enforced a minimum of 1 neighbor per node (min-k = 1). The model used 3 hidden layers with hidden and embedding dimensions of 256, and dropout 0.1. Moreover, PLM-based models and the biophysical feature–based ensemble exhibit similar overall performance on the IgG dataset, with the ensemble showing a slight edge on several metrics (e.g., ROC, AUPRC, MCC).

Cross-validation results (Suppl. Table 8) further clarify these trends. For VHH, the PLM baseline attains the best mean accuracy (0.86), precision (0.83), ROC-AUC (0.87), AUPRC (0.81), and MCC (0.62), while PLM-Disentangled achieves the best mean recall (0.82) and mean F1 (0.74). For IgG, PLM-GNN-Disentangled achieves the strongest mean performance overall, including accuracy (0.68), precision (0.54), F1 (0.64), AUPRC (0.60), and MCC (0.41), whereas PLM-Disentangled yields the highest mean recall (0.86). Together, these results suggest that disentangled attention tends to increase sensitivity (recall), while structure-aware modeling provides the most consistent gains in overall performance for IgG, and does not improve mean VHH performance under the current cross-validation setting.

### 3.5 Explainability of models predictions

Next, we examine the key factors that drive the model predictions for both the biophysical feature–based and transformer-based approaches.

#### 3.5.1 Model-derived biophysical drivers of self-association

One of the key advantages of biophysical feature-based models is the interpretability of their outputs. We leverage this property to investigate the biophysical substrates underlying elevated CSI-BLI for VHHs. We focus on VHHs for this analysis due to better model performance. We use the widely adopted SHAP package to quantify the contributions of individual features to model predictions. We focus on the Gradient Boosted Tree component of the Ensemble, which enables fast and accurate SHAP computation for tree-based models. A beeswarm plot of the resulting SHAP values is shown in Fig. 3a. Features related to aggregation, hydrophobicity, and charge contribute strongly to predictions: in particular, a higher overall dipole moment and increased aggregation propensity within the CDRs are associated with elevated self-association risk, whereas aggregation propensity in the framework region (as measured by Aggrescan (Conchillo-Solé et al. 2007)) is associated with reduced risk.

To further elucidate interactions among these features, we employ a *supervised* clustering approach in SHAP space: we cluster positive training samples (Class I) according to their SHAP value profiles (see Methods for details). This analysis reveals distinct clusters that correspond to different biophysical drivers of self-association, as shown in Fig. 3b-c. Cluster 1 is predominantly characterized by charge-related features (strong dipole moment and formal charge in the CDR region), whereas Cluster 2 and 4 exhibit pronounced aggregation/hydrophobicity related components, either in the CDR or in the framework region. Cluster 3 shows a mix of charge and aggregation components, having both strong dipole moment and formal charge and low AggreScan in the framework. These findings align with prior suggestions that both hydrophobicity and charge are important determinants of self-association and non-specific binding (Gupta et al. 2022; Makowski et al. 2024).

Notably, model performance varies across clusters: sequences with both strong charge and aggregation signatures from Cluster 3 are predicted more accurately, while performance is lower for Clusters 1, 2 and 4, see Fig. 8. This pattern is consistent with the observation that Cluster 3 sequences exhibit higher CSI-BLI, Fig. 8.

#### 3.5.2 Disentangled Attention Component Analysis

We quantify the contribution of each disentangled attention component using the *mean absolute attention score*. For each test sample *b* and component *c* ∈ {CC, CP, PC, CS, SC}, we obtain a head-averaged component score matrix 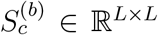, where *L* is the number of residues. We define the per-sample mean absolute attention score as

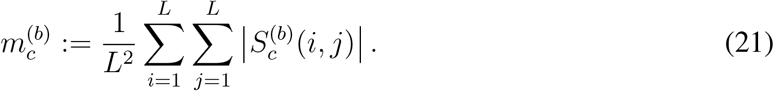

The absolute value ensures 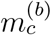 captures the overall *magnitude* (unsigned contribution) of component *c* aggregated over all residue pairs. To summarize relative component usage, we convert these scores into per-sample proportions:

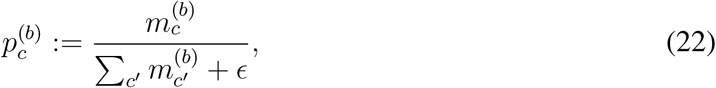

and report the test-set mean 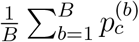 for each component (visualized as a pie chart).

Figure 4 compares the component distribution of the PLM-GNN-Disentangled model (Fig. 4a) to the PLM-Disentangled model (Fig. 4b). For the PLM-Disentangled model, the attention mass is mostly concentrated in the positional interaction channels (CP and PC), while CC contributes little. A plausible explanation is that PLM embeddings (ESM) already encode substantial content-based residue dependencies through their internal self-attention, making an additional content-content pathway in the downstream disentangled block less necessary. In this setting, the module primarily uses positional pathways to refine PLM-derived representations.

**Figure 4:**
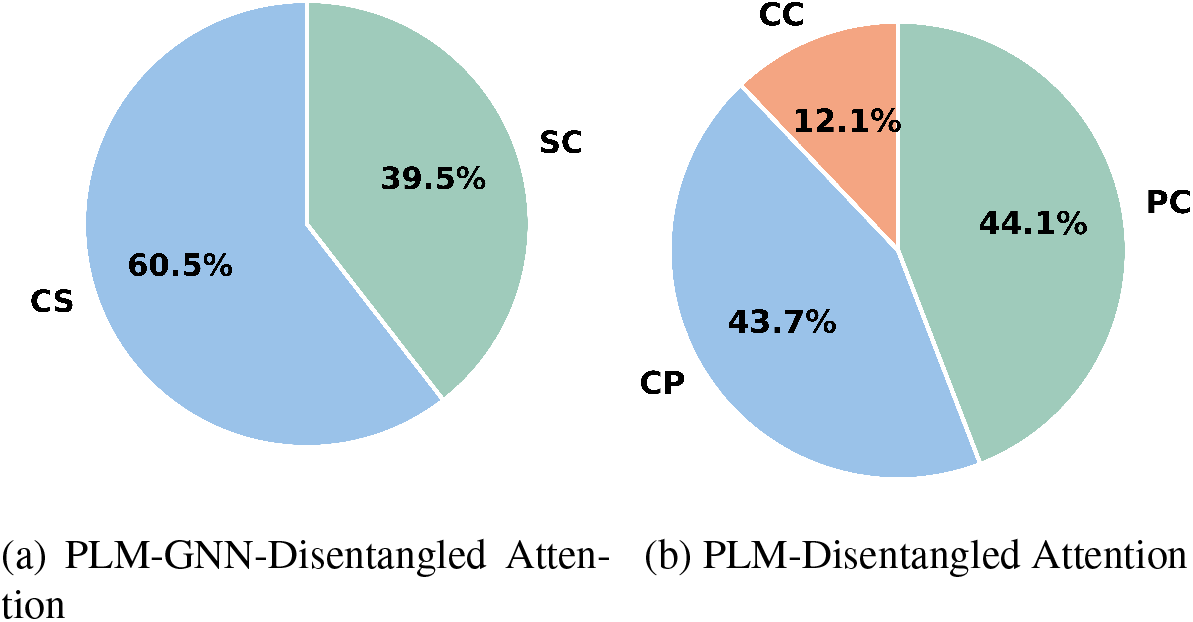
Disentangled attention component distribution on the test set: (a) PLM-GNN-Disentangled, (b) PLM-Disentangled.

**Figure 5:**
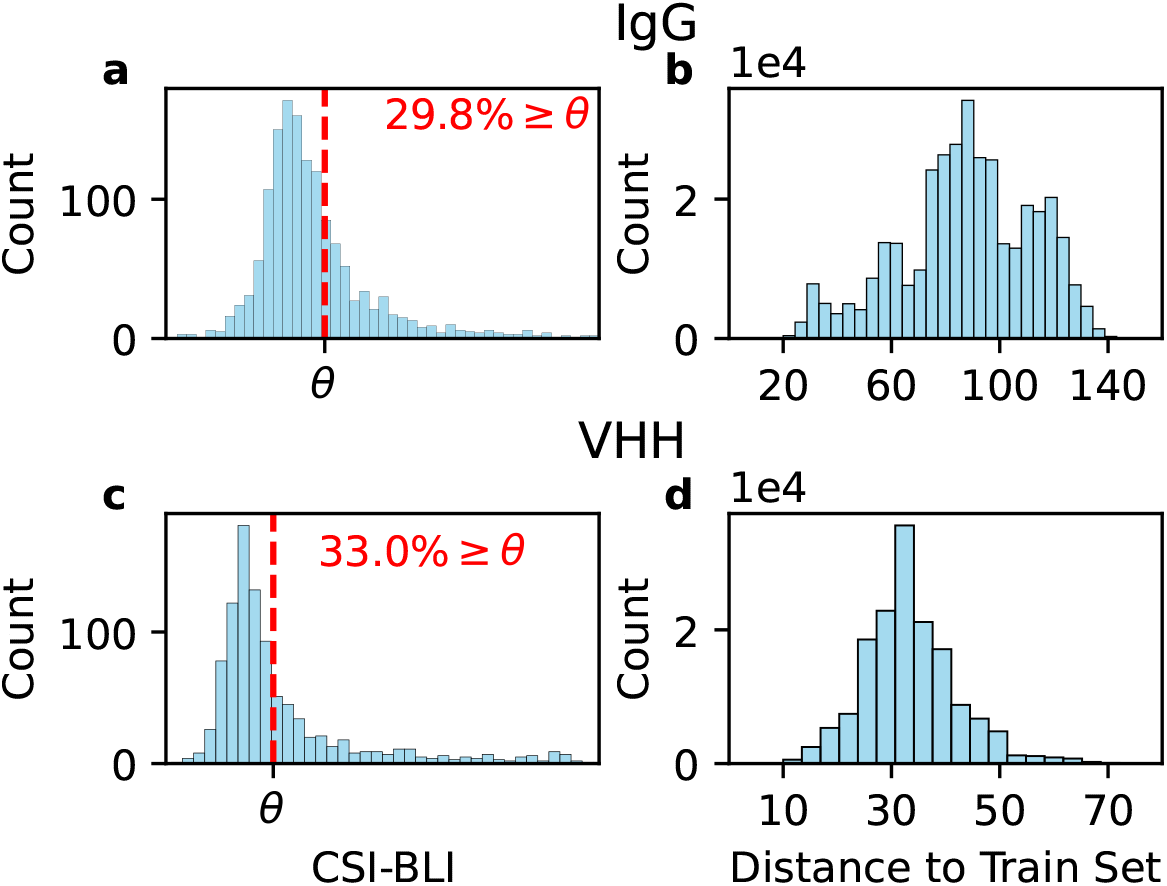
CSI-BLI distribution and train-test sequences pairwise Levhenstein distances for IgG (**a**,**b**) and VHH (**c**,**d**) respectively.

**Figure 6:**
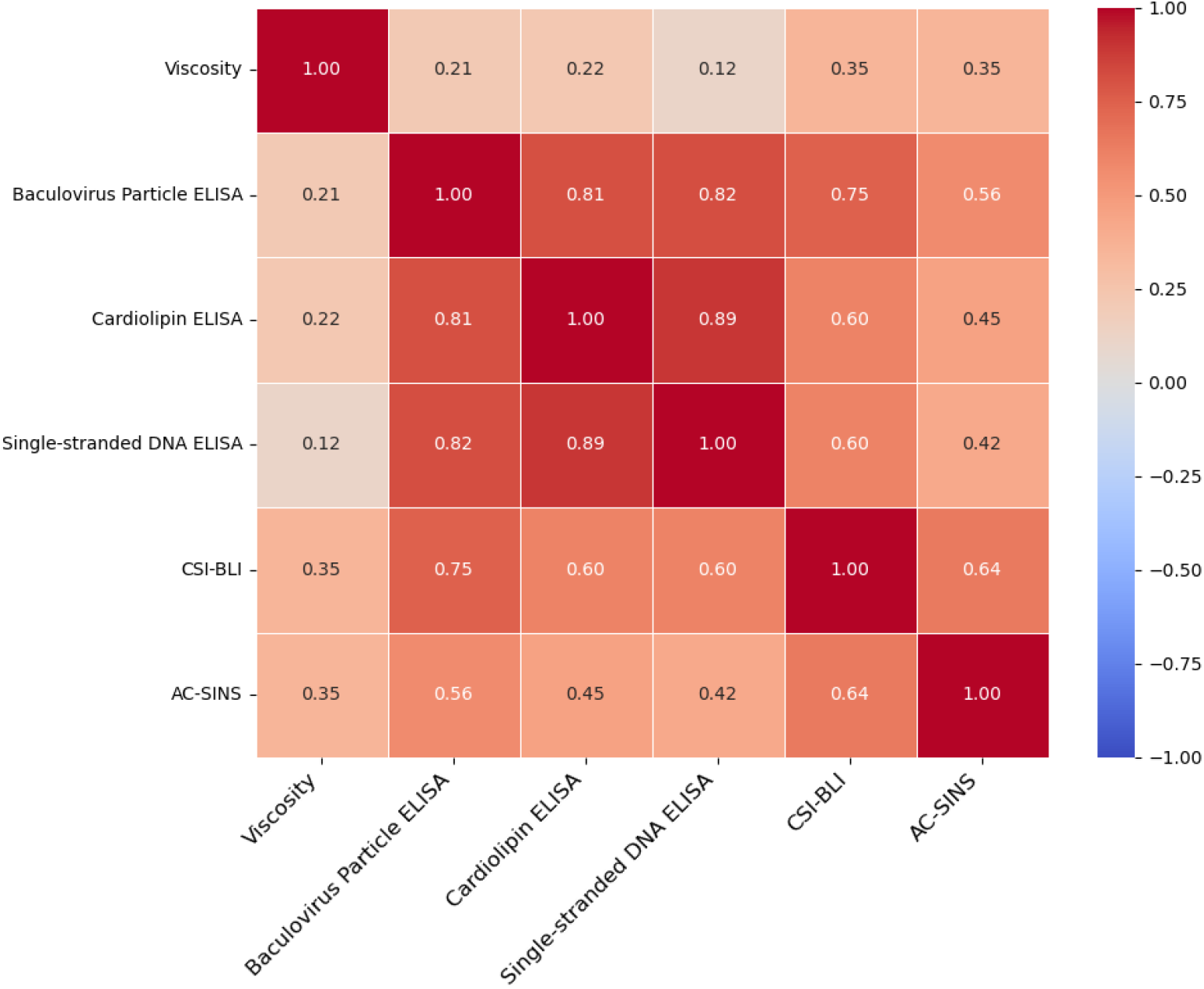
Spearman rank analysis on a 246 mAb dataset shows a moderate positive correlation between viscosity and both CSI-BLI and AC-SINS.

When structural context is introduced via the GNN stream, the attention mass shifts toward cross-modal pathways, with CS and SC dominating the distribution (Fig. 4a). This suggests that, when structure is available, the model preferentially routes information through content-structure and structure-content interactions, using geometry to modulate sequence-derived content representations.

Despite this redistribution toward structural channels, the hold-out results in Table 3 show only modest gains in F1 when adding structure. For VHH, PLM-GNN-Disentangled improves F1 from 0.7451 to 0.7586, and for IgG from 0.5503 to 0.5727. Overall, the component analysis shows a clear shift in how the model uses the disentangled pathways: adding the structural stream moves attention mass from the positional channels (CP/PC) to the cross-modal channels (CS/SC). Yet the hold-out metrics change only slightly, indicating that the structural branch largely reinforces patterns already represented by the PLM and positional pathways rather than introducing a new, dominant source of predictive signal.

Together, the two explainability analyses provide complementary views of what drives CSI-BLI predictions. For biophysical feature models, SHAP yields direct, human-interpretable attributions in terms of physicochemical mechanisms (e.g., charge/dipole, hydrophobicity, aggregation propensity) and reveals distinct subclasses of self-associating antibodies via clustering in explanation space. For transformer-based models, interpretability is more architectural and behavioral than mechanistic: the disentangled attention component analysis clarifies which interaction pathways the model relies on (positional vs. cross-modal content–structure routing) and shows that adding structure primarily reallocates attention mass toward cross-modal channels, even when aggregate predictive performance improves only modestly. Taken together, these results suggest that feature-based models are better suited for mechanistic hypothesis generation, whereas transformer explainability is currently most useful for diagnosing model reliance on modalities and inductive biases rather than pinpointing specific biophysical causes.

## 4 Conclusion

CSI-BLI is a practical, plate-based, automation-compatible assay that directly monitors weak, reversible antibody self-association with minimal material, and it concords with self- and cross-interaction chromatography as well as high-concentration aggregation (Sun et al. 2013). Beyond its established utility for early screening, our analyses position CSI-BLI as an anchor assay informative for downstream liabilities: it associates with formulation viscosity and complements non-specific binding readouts to predict non-target-mediated clearance in hFcRn Tg32 mice, enabling efficient, high-throughput risk triage before resource-intensive viscosity measurements or in vivo PK studies (Jain et al. 2017; Armstrong et al. 2024; Jain et al. 2024). We therefore treat CSI-BLI not only as an experimental benchmark but also as a meaningful modeling target to reduce wet-lab burden through in silico pre-screening and prioritization.

Across antibody formats, predicting CSI-BLI from sequence and structure is feasible but varies in difficulty. Classification is more challenging for IgG than for VHH, reflecting paired-chain complexity and heterogeneous self-association mechanisms in full-length antibodies, consistent with broader developability observations (Jain et al. 2017; Zhang et al. 2023). This difference appears in lower discrimination under edit-distance–controlled IgG hold-out evaluation relative to VHH, emphasizing the need for architectural and data-splitting rigor to avoid sequence-related leakage.

Methodologically, modern transformer-based approaches have comparable or better performance than biophysical descriptor models. Within our architecture, disentangled multi-stream attention yields consistent gains over a standard PLM baseline by explicitly modeling interactions among sequence content, positional context, and structural signals; the sequence–structure fusion variant further improves performance on the harder IgG setting. This supports explicit content–position–structure coupling over feature concatenation or sequence-only attention (Li et al. 2024; Rives et al. 2021; Olsen, Moal, and Deane 2022). Complementarily, biophysical descriptor models constructed from AlphaFold structures and Schrödinger-derived features—paired with cluster-aware selection—offer robust performance and interpretability. Ensembling heterogeneous learners increases generalization, and SHAP analyses reveal mechanistic substrates of elevated CSI-BLI, including charge/dipole, hydrophobicity, and aggregation propensity distributed across CDRs and frameworks (Conchillo-Solé et al. 2007; Lundberg et al. 2020; Makowski et al. 2024). Such mechanistic insights remain harder to extract reliably from deep models without additional analysis, underscoring the value of maintaining both interpretable and high-capacity modeling tracks.

Overall, CSI-BLI is a high-value, high-throughput assay for early developability screening that carries information relevant to viscosity and clearance risks (Sun et al. 2013; Jain et al. 2024; Armstrong et al. 2024). While self-association is only partially determined by sequence and predicted structure, our results show that in silico models—both interaction-aware sequence–structure fusion and interpretable biophysical descriptors—can recover a meaningful fraction of CSI-BLI signal. These models provide practical pre-screens to triage large antibody libraries, prioritize candidates for engineering, and focus experimental validation on the most promising or riskiest molecules, reducing cost and time across discovery-to-development workflows (Bailly et al. 2020; Zhang et al. 2023).

Recommended validation and future directions include expanding sequence-diverse cohorts with standardized CSI-BLI protocols, integrating orthogonal NSB and viscosity assays for multi-task learning, assessing sensitivity to AlphaFold structural uncertainty and surface descriptor parameterization (Park and Izadi 2024), and extending the modular architecture to other endpoints such as polyspecificity and solubility (Yu et al. 2024; Zhou et al. 2025; Sormanni et al. 2017).

## 5 Supplementary information

### 5.1 Data distribution

### 5.2 CSI-BLI as a High-Throughput Predictor of Viscosity

Figure 5.2 shows the correlation between non-specific binding, self-association and Viscosity assays on the 246 mAb dataset. Table 5.2 reports the LDA model performance predicting viscosity for different assays combination.

### 5.3 Biophysical features model

As initial screening, this is the full list of candidate models that were explored for both IgG and VHH CSI-BLI predictions: Gaussian Naive Bayes, Logistic Regression, Linear Discriminant Analysis, Random Forest classifier, Support Vector Machine, Gradient Boosting Classifier. Models were trained for different subset from 2 to all available features. The best subset was chosen as explained in the Methods section.

#### 5.3.1 Shap analysis

Fig. 7 shows clustering performance metrics for the SHAP analysis, while Fig. 8 shows model accuracy on the clusters found in section 3.5.1.

**Figure 7:**
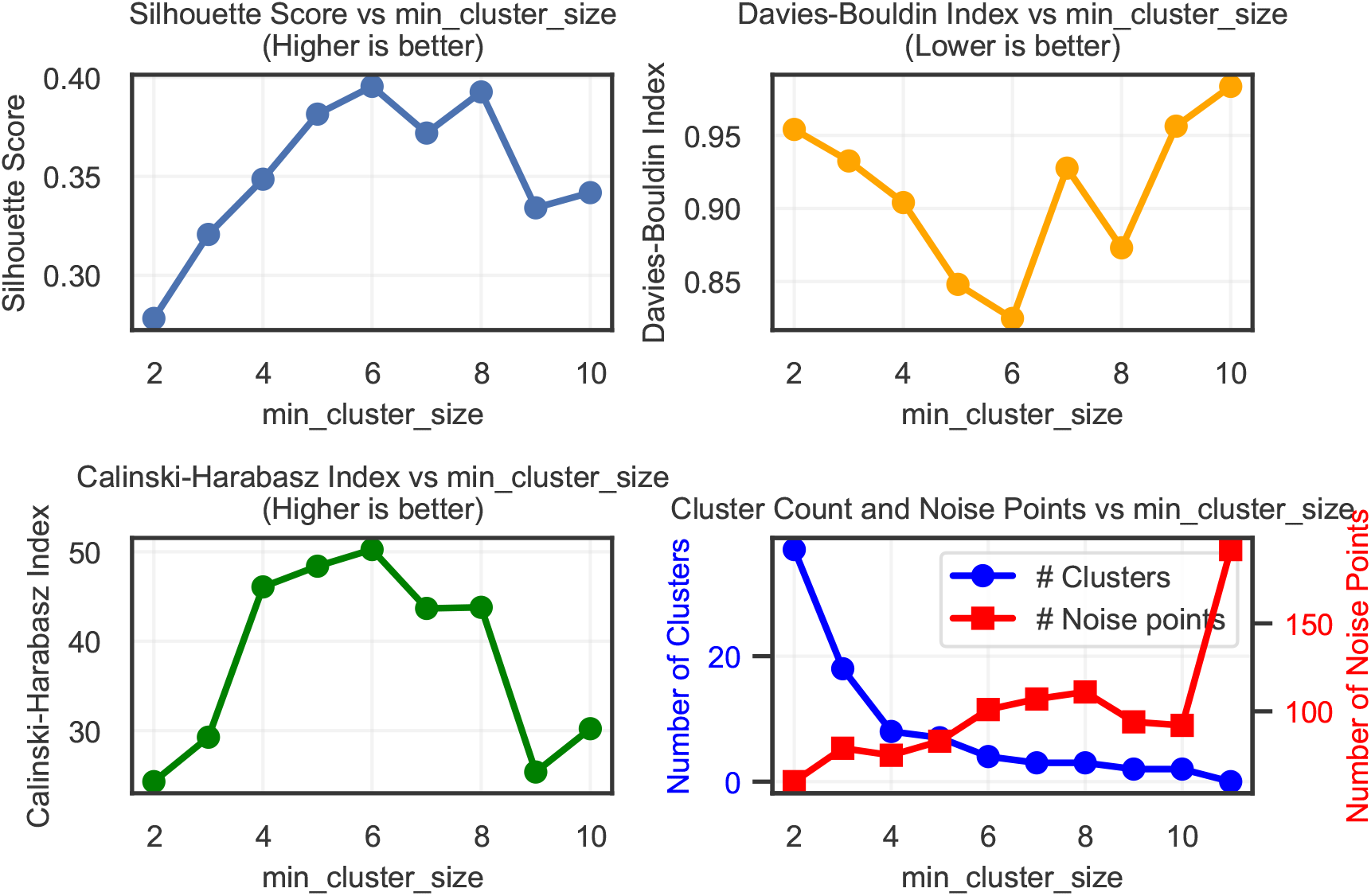
Clustering metrics for increasing values of the minimal cluster size for the HDBSCAN algorithm on SHAP values described in 3.5.1.

**Figure 8:**
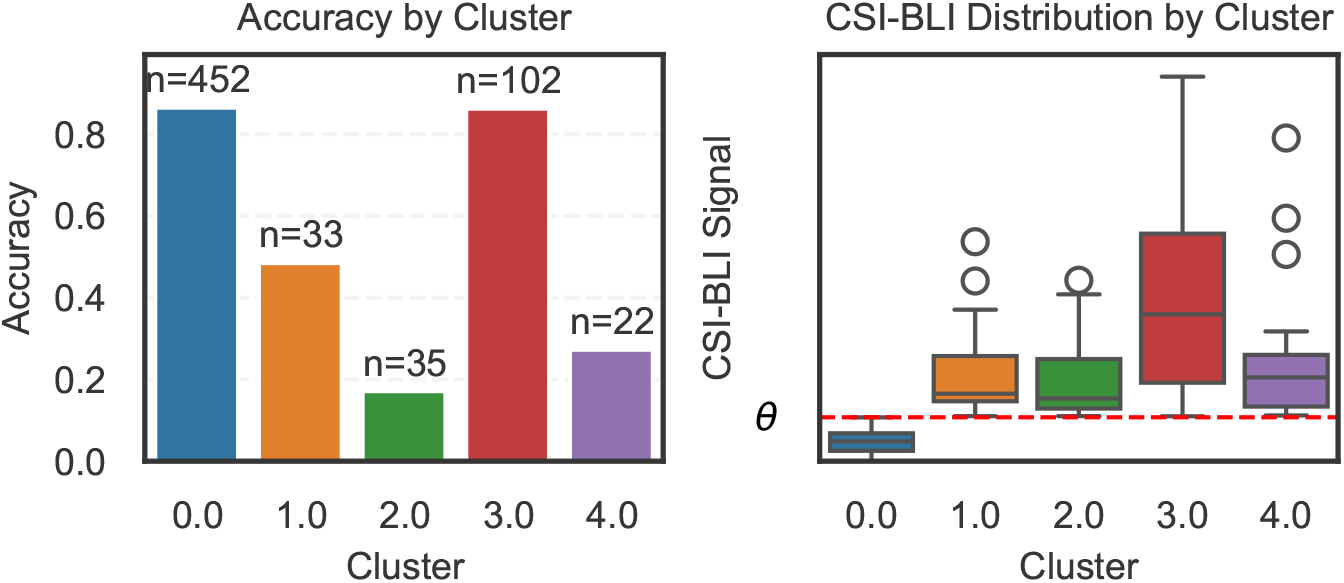
Accuracy and CSI-BLI distribution across the different clusters found in section 3.5.1

**Figure 9:**
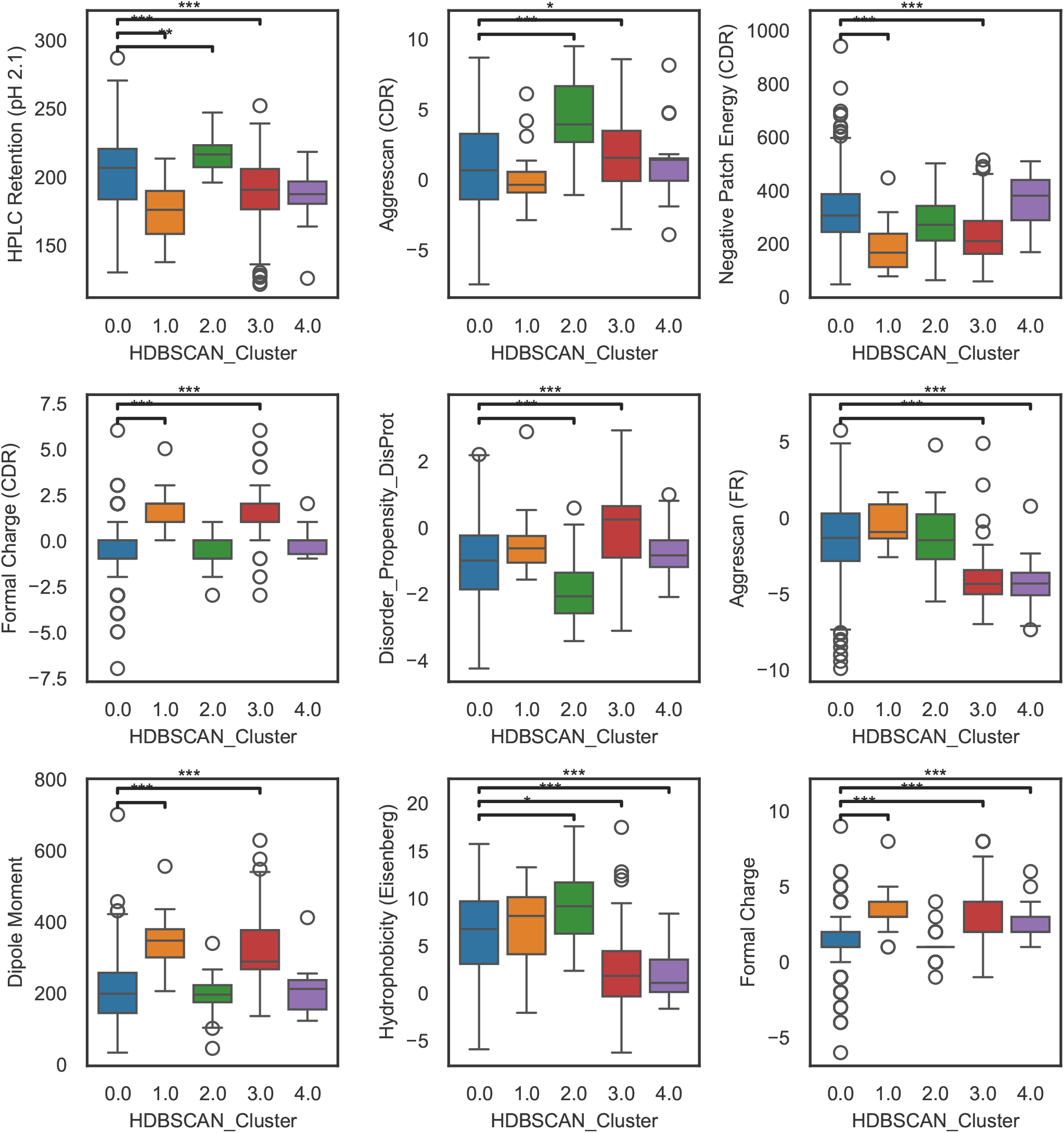
Boxplot of features distributed significantly different across clusters.

### 5.4 Graph Node Features

We follow a standard backbone-geometry parameterization and construct per-residue scalar and vector features from the N–C_*α*_–C backbone atoms.

- **Backbone dihedrals (scalar)**. We compute backbone torsion angles (*ϕ, ψ, ω*) from consecutive N–C_*α*_–C triplets and represent each angle using its sine and cosine. For residue *i*, this yields a 6-dimensional scalar feature:

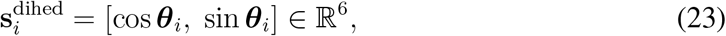

where ***θ***_*i*_ ∈ ℝ^3^ contains the three backbone dihedral angles.
- **Orientation and side-chain proxy (vector)**. Let 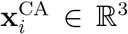 denote the C_*α*_ coordinate of residue *i*. We construct backbone orientation vectors

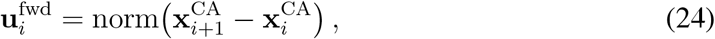

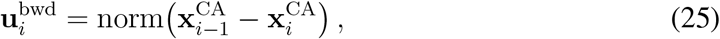

with appropriate padding at sequence boundaries, where norm(·) denotes *ℓ*_2_ normalization. To obtain a side-chain direction proxy, we use the local backbone plane defined by the N–C_*α*_ and C–C_*α*_ vectors. Let

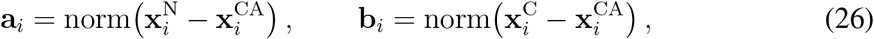

where 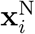 and 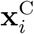 are the backbone N and C coordinates of residue *i*. We compute the in-plane bisector and the plane normal:

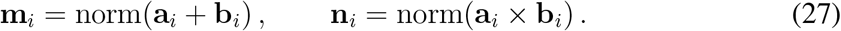

Finally, we combine these two directions into a single unit vector by a normalized sum:

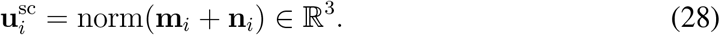

Stacking these vectors yields the initial vector-valued node feature:

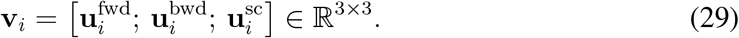

We use the dihedral encoding as the initial scalar node feature:

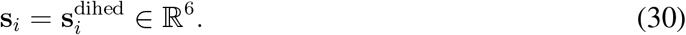

### Edge Features

For each edge (*i, j*), we compute:

- **Distance RBF features (scalar)**. The C_*α*_ distance 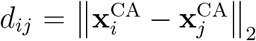 is expanded using a radial basis function (RBF) encoding:

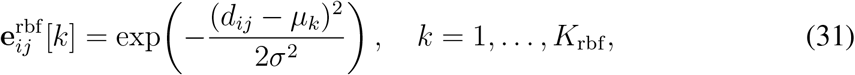

where {*µ*_*k*_} are evenly spaced centers over a fixed interval and *σ* is the bin width.
- **Relative positional encoding (scalar)**. We encode the sequence index difference Δ_*ij*_ = *i* − *j* using sinusoidal embeddings:

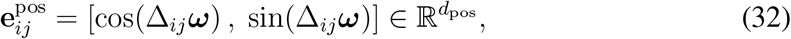

where ***ω*** is a set of frequencies as in Transformer positional encodings.
- **Direction vector (vector)**. We use the unit direction vector from residue *j* to residue *i*,

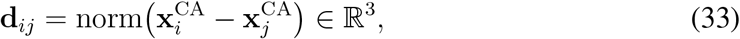

stored as a single-channel vector edge feature.

We concatenate scalar edge features

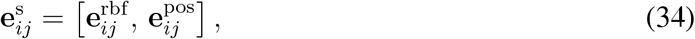

and set the vector edge feature to

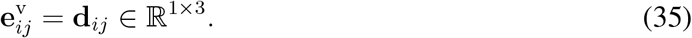

### 5.5 Transformer-based cross-validation results

In table 8 we report the cross-validation results for the transfomer-based architectures.

## 7 Author contributions statement

**Conceptualization**: M.P., S.A., F.D., G.K., N.M.

**Methodology (computational and analytical)**: S.A., F.D., L.L., M.P.

**Experimentation & Data Collection**: T.P., B.A., M.K., A.H., A.L., M.H., C.C., J.D., S.F, A.D.

**Data Processing & Curation**: S.A., F.D., L.L., T.P., B.A., M.K., A.H., A.D., M.P.

**Visualization**: S.A., F.D., M.P.

**Supervision**: M.P., G.K., N.M.

**Writing-Original Draft**: M.P., S.A., F.D., L.L., T.P., A.D.

**Writing-Review & Editing**: S.A., F.D., L.L., T.P., B.A.,A.L., M.H., C.C., J.D., M.K., S.F., A.H., N.M., A.D., G.K., M.P.

## 8 Acknowledgments

We would like to express our sincere gratitude to everyone who contributed to this research. We extend special thanks to Jenna Caldwell, Mehdi Boroumand, Valentin Stanev, Isabelle Sermadiras, Rebecca Croasdale-Wood, and James Savery for their support and facilitation of this work. Assistance from Generative AI tools (OpenAI GPT-5.2 and Anthropic Claude Sonnet 4.5/4.6) was used for language review and code generation for analysis/figure preparation. All content was reviewed, verified, and edited by the authors, who take full responsibility for the accuracy and integrity of the work.

